# The usefulness of triploid sib records for selection of diploid candidates in rainbow trout breeding programs for growth, processing traits and hypoxia tolerance

**DOI:** 10.64898/2026.10.18.759899

**Authors:** Julien Roche, Lou Buisson, Pierre Patrice, Yoannah François, Alexandre Desgranges, Jonathan D’Ambrosio, Pierrick Haffray, Florence Phocas

## Abstract

This study investigates the benefits for rainbow trout selective breeding of using triploid performances rather than diploid ones. Its objectives were to estimate genetic parameters, assess genetic correlations between ploidy levels, and evaluate the gain in selection efficiency using triploid sib testing. The traits analysed were growth, carcass traits, fat content, and acute hypoxia tolerance using pedigree-based and genomic models. Results showed that diploids were heavier and longer than triploids at slaughter age. Triploids exhibited greater tolerance to acute hypoxia in this experiment, although the underlying mechanisms remain unclear.

However, heritability estimates for most traits were similar between ploidy levels when genetic models correctly accounted for triploidy and the unequal maternal and paternal chromosome contributions. Incorrectly treating triploids as diploids substantially overestimated genetic variance and heritability values. Genetic correlations between diploid and triploid performances were high for all traits but varied from 0.60 to 0.98. Correlations were particularly high for carcass yield, fat-related traits and hypoxia tolerance indicating that selection based on diploid performance can efficiently improve these characteristics in triploid offspring. Conversely, correlations for body weight and length were lower, suggesting that evaluating triploid relatives could improve selection accuracy for growth.

Overall, the findings support diploid-based selection as an effective strategy for improving triploid production traits, particularly carcass yield and flesh quality. Direct evaluation of triploid relatives may provide additional benefits for growth-related traits. The study highlights the importance of appropriate genetic and genomic models and recommends assessing the economic value of triploid performance testing for each target trait.

**Highlights:**

- Diploids were heavier than triploids at slaughter age for large fillet production
- Triploids exhibited greater tolerance to acute hypoxia than diploids in this study
- Heritability estimates are similar across ploidy levels for all production traits
- Genetic correlations between ploidy levels are above 0.80 for all traits except growth
- Diploid-based selection is efficient for improving triploid processing traits, but less for growth

## 1. Introduction

Triploid refers to an organism or cell having three sets of homologous chromosomes, i.e. one additional chromosome sets with respect to the number most frequently found in nature for a given species. In animals, triploids are naturally occurring in amphibians, reptiles and fishes (reviewed by Piferrer et al., 2009). For instance, in Atlantic salmon, the occurrence in wild populations is low (Jørgensen et al., 2018), but is higher in farmed populations (Glover et al., 2015). Generally, spontaneous triploids are expected to be sterile due to interference with gametogenesis, resulting in vestigial or highly delayed gonadal development, or to be at least infertile due to random segregation of trivalents producing aneuploid gametes incapable of fertilization. Accordingly, triploids would be regarded as dead ends in reproductive lineages. This, however, is not always the case. In shellfishes, triploids may be produced by the crossing of tetraploid sires obtained from fertile triploids with diploid dams. Several natural diploid– polyploid hybrid complexes are also known in fish, such as the Prussian carp, in which allotriploid females are fertile and reproduce gynogenetically from triploid eggs (Piferrer et al., 2009).

Induction of triploidy by temperature or pressure treatment is a widely practiced and recommended method for producing sterile fish in aquaculture and fisheries management. Induced triploidy is used in various farmed fish and shellfish species to limit immunodepression associated to gonadal maturation and to improve growth, yields and flesh quality in commercial productions (Piferrer et al., 2009). Sterility of triploids can also be used to protect the intellectual property resulting from selective breeding of commercial stocks, as well as to prevent the establishment of invasive species and the genetic introgression of farmed genotypes into wild native populations, in particular for salmonids like *Oncorhynchus mykiss* (Johnson et al., 2019) or *Salmo salar* (Harvey et al., 2017).

To produce triploid fish, shortly after fertilization, eggs are subjected to a temperature or pressure treatment (Piferrer et al., 2009) to inhibit the second meiotic division. This treatment prevents the extrusion of the second polar body from the fertilized egg. This second polar body contains a set of maternal chromosomes and its retention through meiosis II results in a cell with one paternal and two maternal sets of chromosomes. All these three sets of chromosomes are replicated in subsequent cell divisions of the developing embryo. Triploid fish have therefore two sets of chromosomes of maternal origin and one set of chromosomes of paternal origin, with a subsequent increase in cell size but a reduction in cell number.

Triploidy is commercially used in particular in Canada, Chile, France, Japan, UK, and USA for various species of salmonids (mainly rainbow trout, Atlantic salmon, Chinook salmon, Coho salmon, and Artic charr), cyprinids (loach, grass carp), and molluscs (mainly Pacific oyster, but also scallops, clams and mussels) (e.g. Piferrer et al., 2009). Triploid females do not experience the shift from somatic growth to gonadal growth in salmonids, thus preventing mobilization of lipids and deterioration of muscle quality (Manor et al., 2012). In consequence, rainbow trout production of large fillets in fresh water conditions and/or under long photoperiodic regimes in recirculated systems is mainly based on all-female stock and triploid fish.

Some negative side effects of triploidization has been frequently reported in salmonids, such as lower growth, higher sensitivity to abiotic stress or higher vertebral or bones malformation rates when environmental conditions become limiting (Benfey and Biron, 2000; Kacem et al., 2004; Fraser et al., 2012). Reducing such side effects by genetic selection on triploid performances may support welfare improvement and societal acceptance of these induced triploid genotypes. Genetic selection for improving triploid performance is of interest but, because triploids are sterile, selective breeding programs must be applied to diploids parents expecting that the genetic gain can be largely transferred to their triploid progeny. However, clear information is lacking to decide if using diploid phenotypes as selection criteria for improving triploid performance is relevant, or if triploid phenotypes are needed for an efficient breeding program. Until now, due to the cost and difficulty to rear in parallel a sufficient number of diploids and triploids families, only Johnson et al. (2007) estimated heritability of growth and survival in chinook salmon, using a 31 paternal half-sib families’ design that allowed the estimation of additive genetic variances for triploids based on sire variance estimates. Using ANOVA approaches in limited experimental designs, significant family × ploidy interactions have been repeatedly observed for fish weighted less than 1 kg in various salmonids (Withler et al., 1995; Choubert et al., 1997; Bonnet et al., 1999; Blanc et al., 2001; Friars et al., 2001; Johnson et al., 2007; Chiasson et al., 2009; Shrimpton et al., 2012; Weber et al., 2014). Most of these previous works suggest that selective breeding programs may benefit from integration in the genetic evaluation of triploid performance, in particular for fork length, weight and growth as well as for flesh color. Significant genetic response in triploids to selective breeding in diploids was, however, reported (Bonnet et al., 2002). Nevertheless, evaluating only diploid performance may be suboptimal if the genetic correlation between diploids and triploids measured for the same trait differs from unity. Hence, maximizing genetic gain in diploid lines requires the evaluation of breeding values on triploid sibs of the diploid candidates. Advances in the development of genomic tools is an opportunity to evaluate a great number of diploid and triploid families in mixed family design. This implies correctly genotyping triploids and recovering their pedigree as previously reported with microsatellites markers in rainbow trout (Haffray et al., 2012) or in turbot (Hernandez-Urcera et al., 2012). The development of methods for analysing autopolyploids has lagged behind those for diploid species (Amadeu et al., 2020; You et al., 2018). However newly developed methods implemented in free-access R packages both for genotype calling (Roche et al., 2024a) and parentage assignment of triploids using SNP markers (Roche et al., 2024b) now pave the way to properly answer these issues for triploids.

To fill the gap of knowledge regarding the indirect responses on hyperbaric triploid commercial performances based on diploid genetic evaluation and selection, the aims of our study were: i) to choose the correct pedigree-BLUP model to estimate genetic parameters when there is a mixture of diploid and triploid information; ii) to compare the estimates of heritability and genetic correlations between diploid and triploid performances for large rainbow trout production traits and tolerance to hypoxia and predict expected selection responses; iii) to assess the usefulness of a genomic evaluation for production traits using a mixture of diploid and triploid genotypes.

## 2. Material and Methods

### 2.1 Ethics Statement

This study used fin clips collected by the breeding company “Milin Nevez” (Bretagne Truite Group, Plouigneau, France) as part of their commercial breeding programs. All challenges were carried out in accordance with the European guidelines (Directive 2010–63-EU) and the corresponding French legislation. Animal experiment procedures were approved by the ethics committee on animal experimentation COMETH ANSES/ENVA/UPC No. 16 and were authorized by the French Ministry of Higher Education, Research, and Innovation under number APAFIS#24441-2020022417122193 v3, project N°20-032.

### 2.2 Establishment and rearing of challenged fish

The experimental design as partly been described in (Prchal et al., 2023). The experimental stock was established in November 2019 from 190 dams and 98 sex-reversed neomales of a commercial selected all-female line of Milin Nevez breeding company using ten independent full-factorial mating designs. From each of the parents used in the mating, a fin clip was collected and stored in 98% ethanol for later DNA extraction and genotyping.

Four batches were produced. Two triploid batches were produced through high-pressure treatment from the 190 spawns, but only the 90 best spawns were used to produce the two diploid batches.

The progeny stock was then reared under common commercial conditions of “Milin Nevez” breeding company (fed ad libitum using extruded commercial feed, non-limiting oxygenation concentration) until 9 months of age.

At 260 dpf, 1508 diploid and 1535 triploid individuals were PIT (passive integrated transponder) tagged by Bretagne Truite. They were transferred to the FORTIOR Genetics platform (SYSAAF-ANSES, Plouzané, France) in two batches (distinct diploid and triploid batches), where they were acclimatized for four days before an acute hypoxia challenge test was performed for the two batches.

The two remaining batches (one diploid, one triploid) of 1500 fish were kept at the Keryas fish farm for rearing under commercial conditions until the age of 19 months, before being slaughtered and cut at the Bretagne Truite production plant (Plouigneau, France).

### 2.3 Phenotype recording

For the first diploid and triploid batches, the fish were challenged to acute hypoxia in August 2020 at average weights of 50.6 g for the diploids and 38.2 g for the triploids. The diploid batch was phenotyped between the 27^th^ of July and the 4^th^ of August 2020, while the triploid batch was phenotyped three weeks later, between the 18^th^ and the 26^th^ of August 2020. The acute challenge to hypoxia has been described in Prchal et al. (2023). Collected phenotypes were weight and time before loss of equilibrium under hypoxic condition (TLE).

The two other diploid and triploid batches were assessed for production traits at slaughter in June 2021. Production traits were assessed for 1500 diploids at 2.31 kg on average and for 1500 triploids at 1.68 kg on average. The diploid batch was slaughtered and phenotyped between the 31^st^ of May 2021 and the 3^rd^ of June 2021, while the triploid batch was slaughtered and phenotyped two weeks later, between the 14^th^ and the 17^th^ of June 2021.

The assessed traits were the body weight (BW) at 19 months, the fork length (FL), the head weight (HeadW), the viscera weight (ViscW), the headless gutted carcass weight (HGCW), the fat content in muscle (MuscleF), the flesh fat content (FleshF) and the subcutaneous fat content in a whole steak (SubcutF). The viscera weight is the amount of intestine, containing organs such as gut, stomach and liver. The headless gutted carcass weight is considered as a proxy of the two fillets weight but with higher heritability (Haffray et al., 2012). Muscle fat content was recorded using a Fish Torry Fat-meter using a microstrip sensor sensitive to water concentration to obtain a calibrated value of the fat concentration in the muscle as described in Douirin et al. (1998). The subcutaneous and the flesh fat contents in a steak were measured by magnetic resonance imaging (MRI) as described in Blay et al. (2021).

### 2.4 Genotyping and parentage assignment

Fin samples from all the 5,466 phenotyped fish and their 256 parents were sent to the INRAE genotyping platform Gentyane (Clermont-Ferrand, France) for DNA extraction and genotyping. The challenged fish were genotyped for 57,501 SNPs using the Rainbow Trout Axiom® 57K SNP array (Palti et al., 2015). For diploids fish, genotypes were acquired with Axiom Analysis Suite software. For triploid individuals, genotypes were acquired using GenoTriplo R package (Roche et al., 2024a).

After quality control (including a minimum call rate of 97% for SNP and sample as well as removing all SNPs with probe polymorphism or multiple locations on the Arlee genome reference assembly (GenBank assembly accession GCA_013265735.3), 24,110 SNPs were kept for further analysis of the production traits and 23,275 SNPs for the analysis of the hypoxia tolerance. Among them, 1000 markers with the highest MAF were used for parentage assignment using the last version of the R package APIS (Griot et al., 2020) that enables parentage assignment of both diploid and triploid offspring (Roche et al., 2024b), and considering a positive assignment error rate set to 2%.

The final datasets included information from 1461 diploid and 1415 triploid genotyped offspring for the analysis of production traits, and from 1291 diploid and 1196 triploid genotyped offspring for the analysis of hypoxia tolerance.

### 2.5. Statistical analyses

#### 2.5.1. Statistical model for production traits

The following bi-trait linear animal model was applied considering the vector **Y_p_** of all the records of a given trait measured on animals of ploidy level p (p=2, or 3):

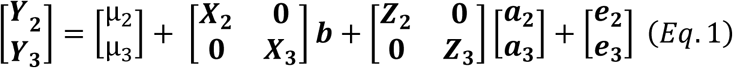

where *µ_p_* is the overall mean performance, ***b*** is the vector of the fixed effect of the testing day (i=1-8) and **X_p_** the corresponding incidence matrix; ***e_p_*** the random residual vector following N(0, **I**σ²_ep_) and 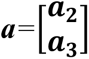 is the random vector of additive genetic effects of animals following either N∼(**0**, **A**⊗***Σ***) for pedigree-based models or N∼(**0**, **G**⊗***Σ***) for a genomic evaluation with:

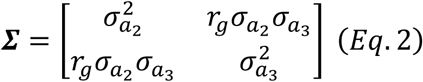

Where σ^2^_ap_ is the genetic variance for animal of ploidy level p and r_g_ is the genetic correlation between the performances of the two ploidy levels. For gutted carcass weight and headless gutted carcass weight, the body weight at slaughter (BW) was added as a covariable in the model and, in consequence, a regression coefficient was estimated in addition to the vector of fixed effects ***b***. For head weight and viscera weight, two different models were run, either adjusting for BW or not, as the adjustment is useful for selective breeding purposes while the raw traits may be more informative from a biological research perspective.

In total, 3473 animals were related through the pedigree relationship matrix (**A**) for BLUP evaluation, tracing back 4 generations of ancestors of the 2876 phenotyped animals for production traits. For the genomic relationship matrix (**G**), only 3126 animals could be related based on the genotypic information of their parents.

#### 2.5.2. Statistical model for hypoxia challenge traits

The following bi-trait linear animal model was applied considering the vector **Y_p_** of all the records of a given trait measured on animals of ploidy level p (p=2, or 3):

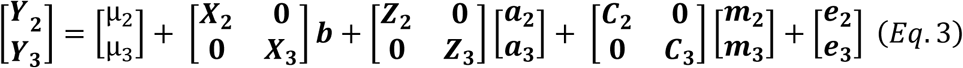

where *µ_p_* is the overall mean performance, ***b*** is the vector of the test day fixed effect (i=1-14) and **X_p_** the corresponding incidence matrix; ***e_p_*** the random residual vector following N(0, **I**σ²_ep_) and 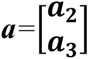 is the random vector of additive genetic effects of animals following N∼(**0**, **A**⊗***Σ***) (see Eq.2 for ***Σ***) ; 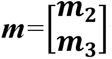 is the vector of the random common maternal effects (j=1-158) following N∼(**0**, **I**⊗**κ**) and 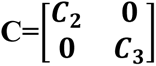 is the corresponding matrix.

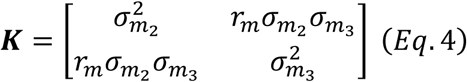

This maternal effect was found significant for hypoxia challenge traits both in diploids (Prchal et al. 2023) and triploids in the current study.

In total, 2975 animals were related through the pedigree relationship matrix (**A**) for BLUP evaluation, tracing back 4 generations of ancestors of the 2487 phenotyped animals for hypoxia. For the genomic relationship matrix (**G**), only 2774 animals were related using the information of their genotyped parents.

#### 2.5.3. Estimation of genetic parameters

Genetic and residual (co)variances were estimated using AIREML algorithm from BLUPF90 software (Misztal et al., 2014). The A-inverse matrix (**A*^−^*^1^**) was given as an input parameter to the algorithm and was generated using polyAinv R package (Hamilton and Kerr, 2018a). The **G** matrix was derived using the new estimator for constructing the genomic relationship matrix **G** proposed by Bilton et al. (2024) in the *R* package GUSrelate for autopolyploid species.

To ensure that the **G** matrix was invertible, a “stabilization” step was necessary. This was achieved by adding a small diagonal component to the **G** matrix as such the transformed matrix was equal to 0.95 **G** + 0.05 **I.**

Analyses were carried out both on single and bi-trait models, considering the two ploidy levels as two different traits. As results were consistent for variance estimates, we chose to focus the presentation of results on bi-traits models. Uni-trait estimates of genetic parameters are, however, given in Supplementary Table 3 for production traits and Supplementary Table 4 for hypoxia challenge traits.

Phenotypic variance (σ²_p_) was calculated for each ploidy level as the sum of the genetic variance and the residual variance for the production traits, to which was added the maternal variance for the hypoxic challenge traits.

Heritability of the diploid and triploid traits (h^2^) were calculated as well as the correlation (r_g_) between the two ploidy expressions of a same trait.

#### 2.5.4. Two ways of modelling triploidy and derivation of the A-inverse matrix

Two genetic models were compared: an “incorrect” and a “correct” one in terms of accounting for triploid animals. The difference between the two models relies on the definition of the **A** and **A*^−^*^1^** matrices. The incorrect model had its **A*^−^*^1^** matrix build as if all phenotyped animals were diploids. Hence, triploids in this “uncorrect” model were considered as diploids. The “correct” model had its **A*^−^*^1^** matrix that considered the right allelic dosage of the triploid offspring, notably the twice higher contribution of the maternal than the paternal genes.

A-inverse matrix was generated through polyAinv R package (Hamilton and Kerr, 2018a) considering the offspring and 4 generations of ancestors. To correctly model the genetic relationships, this package accounts for the probability (λ) that two gametic genes, drawn at random from a randomly selected locus, are inherited from a single parental chromosome (Hamilton and Kerr, 2018a). The mechanism by which unreduced gametes are generated can affect the proportion of gametic genes that are identical by descent, which must be accounted for in the computation of **A*^−^*^1^**. The primary cytological process that leads to the formation of unreduced gametes in diploid parents is the retention of the second polar body by suppressing telophase II of the meiosis in salmonids (Piferrer et al., 2009), which corresponds to a second division restitution (SDR) in plants and the retention of both sets of sister chromatids during Meiosis II (Bretagnolle and Thompson, 1995). In the absence of crossing over, this SDR results in the two genes at some loci being identical by descent as they are inherited from a single parental chromosome. In the case of diploid parents, Hamilton and Kerr (2018) indicated that the parameter λ can be expressed as a function of *β,* the average probability of an odd number of crossovers between the centromere and loci influencing the trait under study. By assuming that these loci are randomly distributed across the genome in the case of SDR, they derived *λ* as equal to 1 – *β* i.e. the probability of absence or of an even number of crossing overs between the centromere and the loci.

To get an approximation of λ that fit our data, we estimated the overall recombination rate *β* of the female gametes. The genotype of parents and offspring and the corresponding pedigree being available, we estimated the recombination rate for a set of 21,047 validated SNPs in high resolution category for both triploid offspring and diploid parents, and for which we identified at least 225 offspring whom the sire was homozygous and the dam was heterozygous. Knowing then the sire’s contribution and the offspring genotype, we could identify the two alleles corresponding to the dam contribution. We counted for a given SNP the number of offspring for which the dam gave twice the same allele (meaning no or an even number of recombination occurred) and the number for which the dam transmitted two different alleles (i.e. an odd number of recombination occurred):

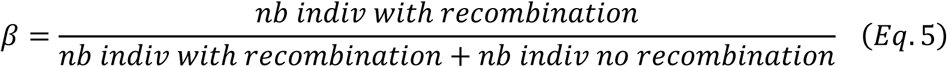

We took the mean of the 21,047 recombination rates and found an overall recombination rate of 0.48. The recombination rate change along the chromosome from 1 (at the telomere) to 0 (at the centromere). Hence, it was expected to get a value close to 0.5, comforting our results. In theory, the recombination rate should be calculated based on the regions of the genome controlling that trait under evaluation. As we expect that all traits under study are highly polygenic with loci influencing the traits randomly distributed across the genome, the mean recombination rate *β* should be meaningful for most traits of interest.

From Eq.4, λ was estimated at 0.52 to derive the **A*^−^*^1^** matrix (Appendix A).

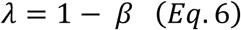

#### 2.5.5. An accurate way to derive genomic relationship matrix for autopolyploids

The new estimator for constructing the genomic relationship matrix **G** proposed by Bilton et al. (2024) aims at improving the estimates of genomic relatedness for autopolyploid species by accounting for errors associated with low-depth high-throughput sequencing data. As, in our case, we had SNP array-based genotype data, we constructed G in a fixed and high SNP read depth setting in order to get reliable estimates of relatedness. As underlined by Bilton et al. (2024), high SNP read depth is an important issue to get unbiased estimates of self-relatedness. Therefore, we assume that all homozygous genotypes corresponded to 240 reads, while heterozygous genotypes corresponded to 120 reads for each allele of a diploid individual, and to 80 or 160 according to the copy number of the reference allele for a triploid heterozygous. By construction, the G matrix accounts for all forms of inbreeding but is unable to distinguish inbreeding due to double reduction from inbreeding resulting from mating related individuals. A noticeable advantage of genomic estimates of relatedness over pedigree-based estimates is that as it automatically captures variation of inbreeding (Figure 1), and therefore double reduction events across the genome based on the genotypic information. Therefore, it bypasses the limitations of estimates from the pedigree-based numerator relationship matrix whose derivation requires fixed genome-wide values of double reduction and recombination rates to be pre-specified (see 2.5.4). In Supplementary Table 1, means and standard deviations of coefficients are given for the two models according to the relationship degree and the ploidy level of phenotyped fish. These statistics showed that all genomic mean estimates of relatedness are slightly lower than their corresponding pedigree-based estimates, but that their variability across fish pairs is tremendously larger.

**Figure 1.**
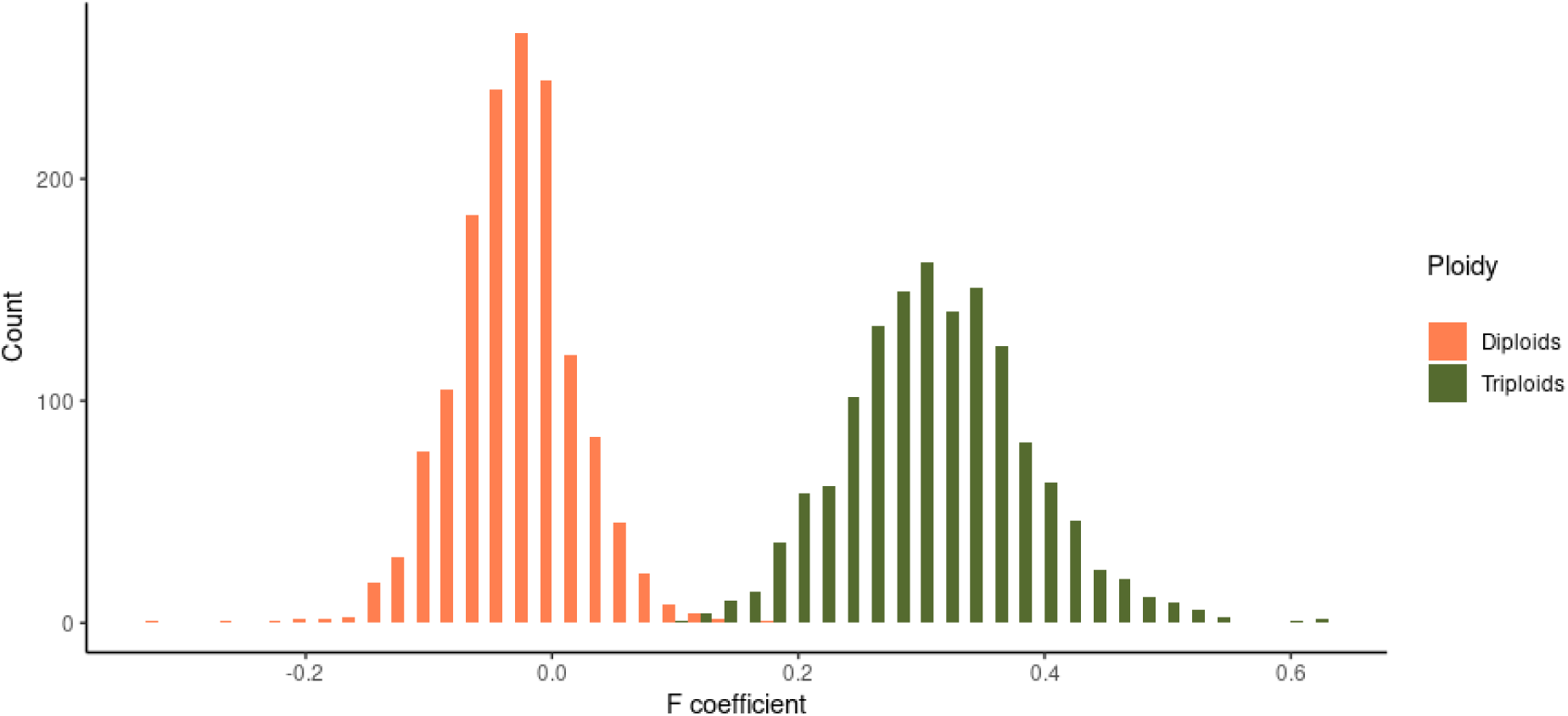
Inbreeding coefficient distribution according to the genomic relationship estimates of the diploid and triploid fish phenotyped for the production traits.

#### 2.5.6. Expected selection responses for triploid expressions of the production traits depending on triploid *vs* diploid collateral tests

In order to assess the interest of implementing a collateral test on triploid production traits, direct and indirect selection responses were derived according to the nature (2n or 3n) of the collateral test implemented in a breeding program.

The direct genetic responses *R_y_* are the responses expected for the triploid phenotypes (***y***) based on selection on triploid sib performances (3n): 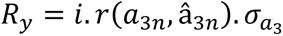, where i is the selection intensity (equal to 1.75 for a selection rate of 10%), *r*(*a*_3*n*_, â_3*n*_) is the accuracy of selection for a collateral test on 3n performances, and 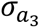 is the additive genetic variance of the triploid expression of the targeted trait ***y***.

The correlated genetic responses are the indirect responses expected for the triploid phenotypes based on selection on performances of diploid sibs (2n): 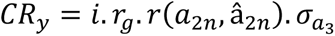, where *r_g_* is the genetic correlation between ploidy expressions of the target trait ***y*** and *r*(*a*_2*n*_, â_2*n*_) is the accuracy of selection for a collateral test on 2n performances.

The technical efficiency of implementing a 3n test in the breeding program rather than a 2n test is derived as the ratio: 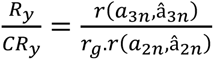

## 3. Results

### 3.1. Raw mean and variance for all recorded traits

Mean performances for most production traits were higher for diploids than triploids (Table 1). Only flesh fat and subcutaneous fate means were slightly higher for triploids, but the differences were not at all statistically significant. For all production traits, CV were very similar (±0.01) between diploid and triploid records.

**Table 1.** Phenotypic mean, standard deviation (SD) and coefficient of variation (CV) for both diploid and triploid performances in production and hypoxia phenotyping tests.

| Tests | Traits | Diploids |  |  | Triploids |  |  |
| --- | --- | --- | --- | --- | --- | --- | --- |
|  |  | Mean | SD | CV | Mean | SD | CV |
| Production | Body Weight (BW, g) | 2327 | 360.9 | 0.16 | 1688 | 278.9 | 0.17 |
|  | Fork Length (FL, mm) | 522.9 | 25.44 | 0.05 | 481.8 | 25.72 | 0.05 |
|  | Head Weight (HeadW, g) | 251.5 | 39.12 | 0.16 | 201.4 | 30.63 | 0.15 |
|  | Viscera Weight (ViscW, g) | 198.1 | 38.64 | 0.20 | 142.3 | 29.66 | 0.21 |
|  | Gutted Carcass Weight (GCW, g) | 2121 | 331.0 | 0.16 | 1539 | 255.1 | 0.17 |
|  | Headless Gutted Carcass Weight (HGCW, g) | 1870 | 297.6 | 0.16 | 1337 | 228.4 | 0.17 |
|  | Muscle Fat (MuscleF, %) | 15.43 | 2.973 | 0.19 | 14.68 | 2.600 | 0.18 |
|  | Flesh Fat (FleshF, %) | 15.25 | 1.881 | 0.12 | 15.65 | 1.708 | 0.11 |
|  | Subcutaneous Fat (SubcutF, %) | 29.61 | 2.977 | 0.10 | 29.64 | 2.839 | 0.10 |
| Hypoxia | Weight (g) | 50.80 | 11.79 | 0.23 | 38.26 | 11.58 | 0.30 |
|  | Time to Loss of Equilibrium (TLE, min) | 190.3 | 72.36 | 0.38 | 217.1 | 61.56 | 0.28 |

Regarding data from the hypoxia challenge batch, the average weight at the challenge time was higher for diploids (50.8 *vs* 38.3 g for triploids) despite the fact that the hypoxia challenge was run three weeks earlier for the diploids. The CV was higher for triploid weight at challenge time compared to the diploids (30% *vs* 23%). On the contrary, the TLE mean was higher for triploids by almost half a raw standard deviation (Table 1), but the CV was lower (28% *vs* 38%) compared to the diploid TLE.

### 3.2. Estimates of variance components and heritability for production traits

#### 3.2.1. Estimates under correct versus incorrect pedigree-based models

Under the correct pedigree-based model, the genetic variances were in general significantly lower for triploid than for diploid traits, except for FL (Table 2). However, the heritability estimates of triploid expressions were not statistically significant different from those of their diploid counterparts, even for SubcutF heritability that exhibited the largest difference in estimates across ploidies (0.42 in triploids *vs* 0.55 in diploids). For all production traits, the genetic coefficients of variation were also quite similar (±0.02) between diploid and triploid expressions.

**Table 2.** Genetic et phenotypic parameters for diploid and triploid expressions of the production traits estimated by the correct ploidy pedigree-based bi-trait model.

|  | Diploid pedigree-based parameters |  |  | Triploid pedigree-based parameters |  |  |
| --- | --- | --- | --- | --- | --- | --- |
| Traits | $\sigma^2_{a2}$ | $\sigma^2_{P3}$ | $h^2$ | $\sigma^2_{a3}$ | $\sigma^2_{P3}$ | $h^2$ |
| *Body Weight (BW, g)* | 32189±7180 | 131630 | 0.24±0.05 | 18497±3426 | 72086 | 0.26±0.04 |
| *Fork Length (FL, mm)* | 115.0±31.47 | 651.9 | 0.18±0.05 | 127.2±26.154 | 624.5 | 0.20±0.04 |
| *Head Weight (HeadW, g)* | 328.3±76.89 | 1513 | 0.22±0.05 | 209.1±39.05 | 863.8 | 0.24±0.04 |
| *Viscera Weight (ViscW, g)* | 672.8±114.2 | 1527 | 0.44±0.06 | 346.3 ±51.63 | 773.9 | 0.45±0.06 |
| *Head Weight (HeadW\*, g)* | 104.3±21.55 | 350.1 | 0.30±0.05 | 60.45±10.07 | 184.1 | 0.32±0.05 |
| *Viscera Weight (ViscW\*, g)* | 403.8±59.37 | 680.8 | 0.59±0.07 | 186.9±23.81 | 282.2 | 0.66±0.06 |
| *Gutted carcass weight (GCW\*, g)* | 410.8±62.25 | 726.2 | 0.57±0.07 | 179.2±23.48 | 293.9 | 0.61±0.06 |
| *Headless GCW (HGCW\*, g)* | 443.4±73.46 | 1022 | 0.43±0.06 | 196.0±28.01 | 431.0 | 0.45±0.05 |
| *Muscle Fat (MuscleF, %)* | 5.431±0.711 | 8.850 | 0.61±0.06 | 3.655±0.451 | 5.989 | 0.61±0.06 |
| *Flesh fat (FleshF, %)* | 2.313 ±0.332 | 3.691 | 0.63±0.07 | 1.496±0.193 | 2.487 | 0.60±0.06 |
| *Subcutaneous fat (SubcutF, %)* | 4.904±0.750 | 8.913 | 0.55±0.06 | 3.091±0.459 | 7.410 | 0.42±0.05 |

\* Corrected for BW

Heritability and variance component estimates of diploid production trait expressions were robust to the change of **A** matrix (correct *vs* incorrect gene flow rules). Phenotypic variances and heritability of diploid expressions were slightly overestimated under the incorrect model (see Supplementary Table 2), differences being under 2% of the correct variance values and less than 8% of the correct heritability values (Table 2). On the contrary, heritability and variance component estimates of triploid expressions were very sensitive to the change of the **A** matrix. Comparing results from the correct and incorrect pedigree-based models (Table 2 vs Supplementary Table 2), we observed severe overestimation of genetic variance and heritability values for triploid expressions using the wrong rules for deriving **A**. When the genetics of triploids was not correctly modelled, genetic variances were overinflated from 66% for SubcutF to up 90% for *HGCW\**, leading to overestimation of both phenotypic variances and heritability values. While overestimation of the phenotypic variances remained moderate (+8 to +27% increases), the heritability of triploid expression was dramatically overestimated under the incorrect model: from values ranging in-between +40 and +45% for the 3 fat-related traits, to values ranging from +55 to +110% for the 4 weight traits with the extreme +110% increase corresponding to BW estimate. Regarding *HGCW*,* the main trait of interest for improving fillet yield, the heritability overestimation was +58% and the phenotypic variance was inflated by 22% (Supplementary Table 2) compared to the correct estimates (Table 2).

#### 3.2.2. Estimates under the genomic model

Estimates of variance components using the **G** matrix (Table 3) were very consistent with those using the correct **A** matrix (Table 2) for all production traits, although a slight underestimation of genetic and phenotypic variances estimated using the pedigree-based models was observed for both ploidy expressions and most of the traits. Absolute differences in heritability values did not generally exceed 0.08, i.e. below 2 standard errors of the estimates. It is to be noticed that the pedigree-based estimates under bi-trait modelling were closer to those of the genomic univariate models considering single ploidy phenotypes (see Supplementary Table 3) than to the genomic bi-trait model combining diploid and triploid performances (Table 3).

**Table 3.** Genetic et phenotypic parameters for diploid and triploid expressions of the production traits estimated by the genomic bi-trait model.

|  | Genome-based diploid parameters |  |  | Genome-based triploid parameters |  |  |
| --- | --- | --- | --- | --- | --- | --- |
| Traits | $\sigma^2_{a2}$ | $\sigma^2_{p2}$ | $h^2$ | $\sigma^2_{a3}$ | $\sigma^2_{p2}$ | $h^2$ |
| *Body Weight (BW, g)* | 30881±5734 | 133940 | 0.23±0.04 | 18872±3022 | 74193 | 0.25±0.04 |
| *Fork Length (FL, mm)* | 143.9±30.05 | 672.1 | 0.21±0.04 | 142.0±24.99 | 644.1 | 0.22±0.04 |
| *Head Weight (HeadW, g)* | 426.6±72.04 | 1569 | 0.27±0.04 | 254.3±38.16 | 897.2 | 0.28±0.04 |
| *Viscera Weight (ViscW, g)* | 725.4±87.02 | 1592 | 0.46±0.04 | 378.5±43.26 | 821.8 | 0.46±0.04 |
| *Head Weight\* (HeadW\*, g)* | 117.1±18.00 | 395.3 | 0.30±0.04 | 55.25±7.792 | 194.4 | 0.28±0.03 |
| *Viscera Weight\* (ViscW\*, g)* | 464.0±42.91 | 723.1 | 0.64±0.03 | 200.5±17.55 | 300.9 | 0.67±0.03 |
| *Gutted carcass weight (GCW\*, g)* | 463.2±45.00 | 766.8 | 0.60±0.04 | 196.4±18.03 | 313.2 | 0.63±0.03 |
| *Headless GCW (HGCW\*, g)* | 555.9±59.44 | 1104 | 0.50±0.04 | 214.8±22.51 | 461.5 | 0.47±0.04 |
| *Muscle Fat*<br>(*MuscleF*, %) | 4.815±0.4844 | 8.869 | 0.54±0.03 | 3.235±0.3171 | 6.161 | 0.53±0.03 |
| *Flesh fat* (*FleshF*, %) | 2.372±0.2198 | 3.792 | 0.63±0.03 | 1.438±0.1341 | 2.636 | 0.55±0.03 |
| *Subcutaneous fat*<br>(*SubcutF*, %) | 4.329±0.4769 | 8.902 | 0.49±0.04 | 3.253±0.3776 | 7.730 | 0.42±0.04 |

\* Corrected for BW

### 3.3. Estimates of variance components and heritability for hypoxia challenge traits

Under the correct pedigree-based model, all the variance components (genetic, maternal and phenotypic variances) were significantly lower for triploid than for diploid traits (Table 4), as it was also observed for genetic and phenotypic variances of production traits (Table 2). While the pedigree-based heritability estimate of Weight was similar across the two ploidies as it has been shown for production traits, the heritability of TLE was estimated at a quite lower value for triploids, though not statistically significantly different, than for their diploid counterparts (0.15 in triploids *vs* 0.28 in diploids). As for all production traits, the genetic coefficient of variation was quite similar (±0.02) between diploid and triploid expressions of Weight. However, in line with the observed difference in heritability estimates of TLE, there was a strong difference in the genetic coefficient of variation for TLE between triploids (9%) and diploids (19%).

**Table 4.**
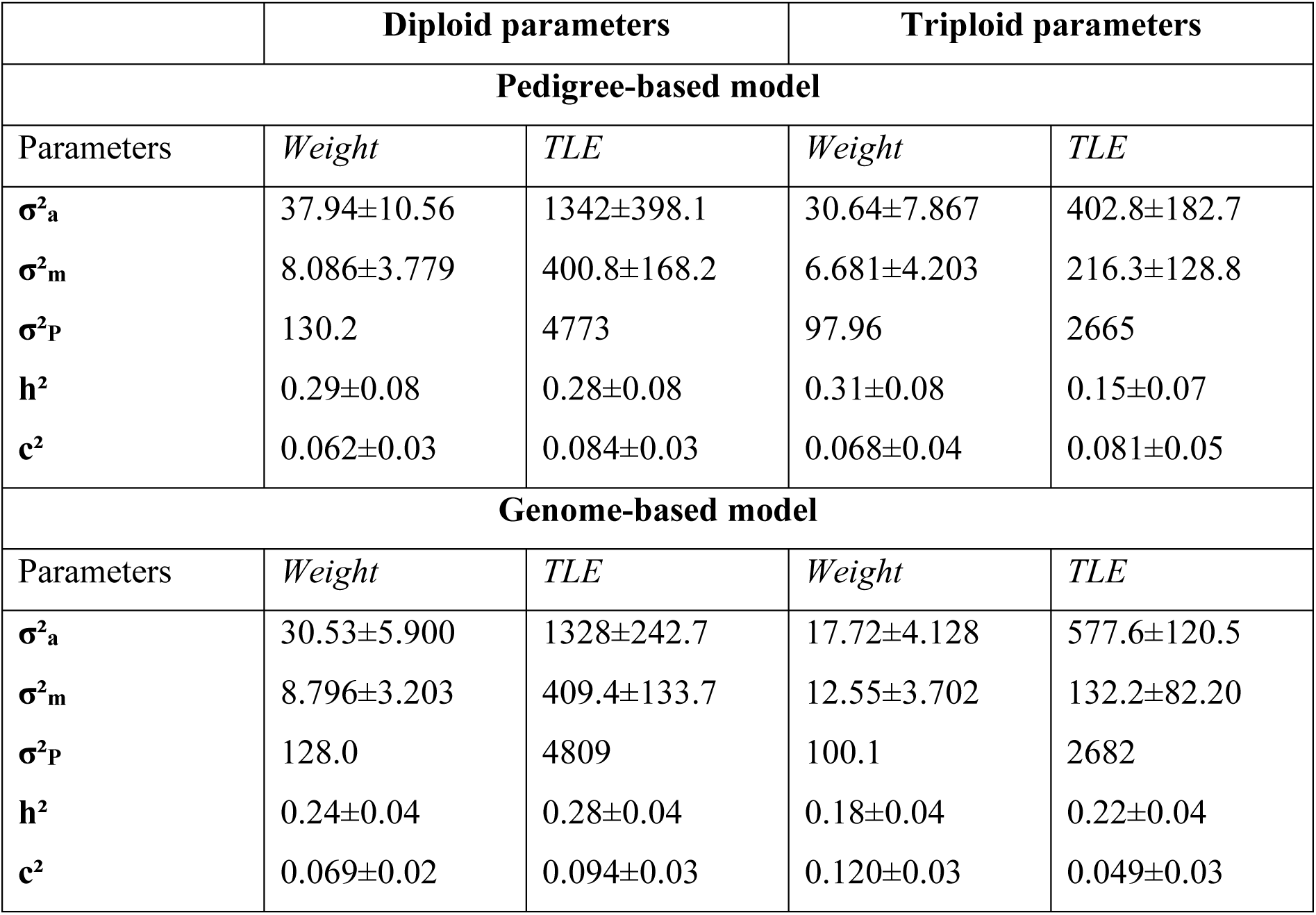
Genetic parameters of the hypoxia challenge traits estimated by bi-trait correct ploidy pedigree and genome-based models.

|  | Diploid parameters |  | Triploid parameters |  |
| --- | --- | --- | --- | --- |
| Pedigree-based model |  |  |  |  |
| Parameters | Weight | TLE | Weight | TLE |
| $\sigma^2_a$ | 37.94±10.56 | 1342±398.1 | 30.64±7.867 | 402.8±182.7 |
| $\sigma^2_m$ | 8.086±3.779 | 400.8±168.2 | 6.681±4.203 | 216.3±128.8 |
| $\sigma^2_P$ | 130.2 | 4773 | 97.96 | 2665 |
| $h^2$ | 0.29±0.08 | 0.28±0.08 | 0.31±0.08 | 0.15±0.07 |
| $c^2$ | 0.062±0.03 | 0.084±0.03 | 0.068±0.04 | 0.081±0.05 |
| Genome-based model |  |  |  |  |
| Parameters | Weight | TLE | Weight | TLE |
| $\sigma^2_a$ | 30.53±5.900 | 1328±242.7 | 17.72±4.128 | 577.6±120.5 |
| $\sigma^2_m$ | 8.796±3.203 | 409.4±133.7 | 12.55±3.702 | 132.2±82.20 |
| $\sigma^2_P$ | 128.0 | 4809 | 100.1 | 2682 |
| $h^2$ | 0.24±0.04 | 0.28±0.04 | 0.18±0.04 | 0.22±0.04 |
| $c^2$ | 0.069±0.02 | 0.094±0.03 | 0.120±0.03 | 0.049±0.03 |

Regarding Weight trait, maternal variance estimate was doubled under the genomic model compared to the pedigree-based one to the detriment of the estimate of genetic variance (Table 4). In consequence, heritability estimate was significantly reduced for W in triploids using a **G**-matrix compared to a **A**-matrix. Conversely, maternal variance estimate for TLE was strongly reduced under the genomic model in favour of an increase in the genetic variance estimate, leading to a higher estimate of heritability of TLE under the genomic model in comparison to the pedigree-based one (Table 4).

This discrepancy in estimates observed between the two models may be due to some confounding issues between genetic additive and maternal effects under the pedigree-based model, a possibility underlined by Hamilton and Kerr (2018). The high uncertainty observed for the variance estimates of maternal effects in both models also illustrates these likely confounding issues. Assuming that the genomic model is the most appropriate to disentangle genetic and maternal effects, we then observed a convergence in the heritability estimates of the diploid and triploid expressions for both Weight and TLE while the proportion of phenotypic variance explained by maternal effects in triploids appeared to be significantly higher for Weight, but significantly lower for TLE than in diploids.

### 3.4. Genetic correlations between diploid and triploid expressions of production traits

All genetic correlations between diploid and triploid performances of production traits were substantial, or even very high (r_g_≥ 0.9), especially for the 3 fat-related traits, as well as for the 4 traits assessing yields through weights of body cuts corrected for whole BW. However, the 3 raw weights (BW, HeadW, ViscW) and FL exhibited correlation estimates ranging from 0.64 to 0.80 that were significantly different from 1 (Table 5). The estimates of genetic correlations between diploid and triploid expressions were very similar under the pedigree and genome-based models accounting properly for the ploidy of all fish (Table 5), with differences smaller than their standard errors. These estimates were also very consistent with those derived under the incorrect pedigree-based model (see Supplementary Table 2), indicating that variance estimates seem more sensitive to the correct ploidy model than correlations.

**Table 5.** Pedigree and genome-based estimates of genetic correlation between diploid and triploid performances for all production traits.

| Traits | Pedigree-based estimate | Genome-based estimate |
| --- | --- | --- |
| *Body Weight (BW, g)* | 0.692±0.132 | 0.638±0.105 |
| *Fork Length (FL, mm)* | 0.793±0.167 | 0.647±0.121 |
| *Head Weight (HeadW, g)* | 0.714±0.133 | 0.684±0.090 |
| *Viscera Weight (ViscW, g)* | 0.794±0.075 | 0.801±0.052 |
| *Head Weight (HeadW\*, g)* | 0.883±0.086 | 0.938±0.053 |
| *Viscera Weight (ViscW\*, g)* | 0.915±0.043 | 0.892±0.031 |
| *Gutted carcass weight (GCW\*, g)* | 0.883±0.050 | 0.886±0.033 |
| *Head gutted carcass weight (HGCW\*, g)* | 0.941±0.053 | 0.974±0.025 |
| *Muscle Fat (MuscleF, %)* | 1.000±0.002 | 0.967±0.028 |
| *Flesh fat (FleshF, %)* | 0.976±0.034 | 0.971±0.023 |
| *Subcutaneous fat (SubcutF, %)* | 0.947±0.051 | 0.883±0.044 |

\* Corrected for BW

### 3.5. Genetic correlations between diploid and triploid expressions of hypoxia challenge traits

As observed for BW, the genetic correlation between diploid and triploid expressions of Weight at the time of the hypoxia challenge was moderate with both pedigree and genome-based estimates about 0.6, significantly different from 1. However, the estimates of the correlation of maternal effects between diploid and triploid Weight were high (r_m_ > 0.9) and almost equal to 1 for the pedigree-based model. Regarding TLE, the genetic correlation estimates between diploid and triploid performances were high and similar across models (r_g_ ∼ 0.9) with lower estimates of the maternal correlation (r_m_ > 0.7). It must, however, be noted the extreme high SE values for all estimates of the correlation of maternal effects between diploid and triploid performances for all traits and models.

### 3.6. Selection accuracy for triploid expressions of production traits according to the ploidy level of the phenotyped sibs and relative efficiency of their direct selection vs indirect selection

Table 7 presents the relative efficiency of direct selection on a triploid sib test **vs** a diploid sib test for all traits, based on the fact that the expected accuracy of a sib test is given by the mean accuracy of the expected breeding values of their parents. Due to differences in the magnitude of the genetic correlation between diploid and triploid expressions across traits, the gain in efficiency may be increased to a greater or lesser extent by using triploid sibs. For instance, Weight, BW and FL would strongly benefit from a selection based on triploid sibs, as their ratio 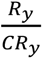 are higher than 1.5. On the contrary, TLE, HGCW*, MuscleF, FleshF and SucutF showed ratio closer to 1, meaning that the use of a triploid sib test would not drastically improve these traits. In particular, ratios below 1 were derived for MuscleF and SubcutF under a pedigree model, indicating that there will even be a slight loss of efficiency using a triploid sib test rather than a diploid one due to lower heritability values for triploids in that case (see Table 2). In addition, genomic selection was only more efficient than pedigree-based selection in using triploid information for BW, FL, MuscleF and SubcutF; conversely, it was less efficient for HGCW*, FleshF, Weight and TLE (Table 7).

**Table 6.** Pedigree and genome-based estimates of genetic correlation between diploid and triploid performances of hypoxia challenge traits.

| Traits | Pedigree-based estimates |  | Genome-based estimates |  |
| --- | --- | --- | --- | --- |
| | $r_g$ | $r_m$ | $r_g$ | $r_m$ |
| *Weight* | 0.614±0.157 | 0.999±0.475 | 0.597±0.135 | 0.908±0.302 |
| *TLE* | 0.901±0.707 | 0.760±0.705 | 0.910±0.095 | 0.868±0.909 |

**Table 7.**
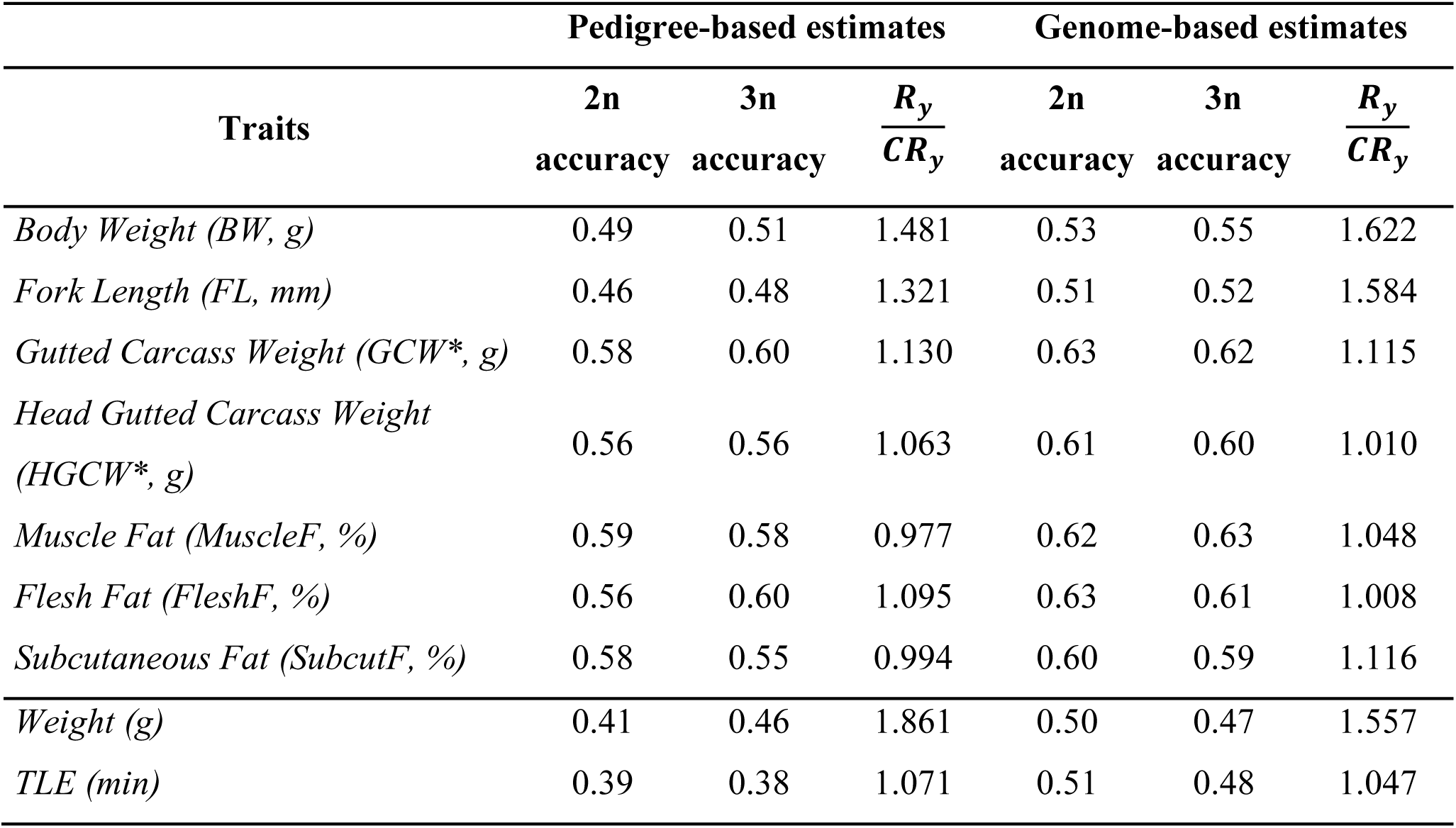
Relative efficiency 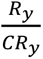 of implementing a triploid sib test rather than a diploid test to improve triploid performances.

## 4. Discussion

It is important to note that triploids discussed here are the results of the retention of second polar body through pressure treatment and not through thermal treatment or the mating of a tetraploid sire and a diploid dam. All conclusions are drawn for the first kind of triploids that is the predominant one nowadays in salmonid production (Blanc et al., 2005). Our results showed that using an incorrect ploidy pedigree-based model for triploid expressions lead to large overestimation of genetic variances and heritability values for triploid performances. Under the correct ploidy model, pedigree-based and genome-based models yielded similar results regarding heritability estimates and phenotypic variances for all production traits. A notable decrease in phenotypic variance was observed in triploids compared to diploids across all models and traits (with the only exception of FL). The heritability estimates of most triploid expressions were quite close to the estimates for the corresponding diploid expressions. Both the phenotypic and genetic coefficients of variation were also quite alike between diploid and triploid production traits. Results were more contrasted for the two traits evaluated in the hypoxia test, probably due to the existence of maternal effects that were difficult to correctly separate from the additive genetic effects. The genetic coefficient of variation in TLE was far much larger for diploids than triploids during hypoxia exposure. However, assuming that the genomic model was the most appropriate to disentangle genetic and maternal effects, we then observed a convergence in the heritability estimates of the diploid and triploid expressions for both Weight and TLE, while the proportions of phenotypic variance explained by maternal effects exhibited opposite trends across the two traits, being significantly lower for Weight, but significantly higher for TLE in diploids than in triploids.

Whatever the traits and models, all genetic correlations between diploid and triploid traits were highly positive, but their estimates for unadjusted weight and length traits showed, however, lower values ranging from 0.60 (Weight at hypoxia test) to 0.80 (Viscera Weight) that were statistically different from 1.

### 4.1. Effect of ploidy on production traits

The performance of triploid sibs of the diploid breeders is of ultimate importance if improved populations must be triploids. In the current study, performances were higher in diploids than triploids both for juvenile weight (at hypoxic challenge time) and for production traits at slaughter age about 19 months. These results are consistent with most studies in the literature of the 80s-90s that showed that, before the age of sexual maturation, reached at 2 or 3 years of age for diploid females, triploidization in salmonids generally induced a survival and growth depression in embryos and larvae with subsequent effects on body weight during a variable length of time (O’Flynn, 1997; Quillet et al., 1988; Utter et al., 1983). Most of this depression is established during the first three months of life, triploids appearing very similar to the control later on. At the end of the immature period (∼50-100g), triploids of rainbow trout exhibit moderate but significant depression relative to the diploid control for both survival (about 25% decrease) and growth (10 to 15% decrease) (Quillet et al., 1988). As reviewed recently by Hansen et al. (2025), it is probable that the generally reduced fitness of triploids is due to the partial inbreeding caused by the doubled maternal genome. We confirmed in our study that triploid offspring are significantly more inbred than their diploid sibs as the mean genomic inbreeding coefficient of triploids was 31.0% compared to -2.6% in average for their diploid counterparts (see Supplementary Table 1). Therefore, the retention of the second polar body in the eggs results in a level of inbreeding in induced triploids that is slightly lower than that expected from two generations of full-sib mating (37.5% of homozygosity in expectation).

However, in a few studies the growth of juvenile triploid salmonids was found to be the same as of diploid counterparts (O’Flynn, 1997; Scott et al., 2015) or event better as mainly reported in juveniles in rainbow trout and brown trout *Salmo trutta* (Blanc et al., 2005). Growth of triploids generally exceeds that of female diploids once gonadal maturation slows the growth of diploids (e.g. Maxime, 2008; Piferrer et al., 2009), at least in freshwater rearing conditions.

This was not true after transfer at 1 year of age to a saltwater tank for over 15 months with diploids then outperforming triploids in growth and survivorship (Johnson et al., 2019).

Fat percentage influences the quality of the final fillet product. Our results for fat-related traits (Table 1) confirmed that the amounts of muscular and visceral lipids are alike in triploids and diploids fishes (e.g.Maxime, 2008), as already shown in rainbow trout (Oliva-Teles and Kaushik, 1990; Sheehan et al., 1999)

For all production traits, CV were alike between diploid and triploid records (Table 1). These results are in contrast with previous studies showing that triploidy was associated with increased CV in growth traits in salmonids (Friars et al., 2001; Johnson et al., 2007; Weber et al., 2014).

### 4.2. Effect of ploidy on acute hypoxia tolerance

Interestingly, triploids exhibited a better tolerance to acute hypoxia exposure compared to their diploid counterparts in our study. To date, most of the physiological research in triploids has focused on the effects of acute stressors such as hypoxia, confinement, handling stress, swimming exercise (Benfey, 1999; Benfey and Biron, 2000; Hyndman et al., 2003; Sadler et al., 2000; Scott et al., 2015). These studies show that the physiological responses are generally very similar between diploid and triploid fish, but none of them investigated long and/or variable hypoxia or other stress regimes that are more representing farming context.

Regarding tolerance to acute hypoxia, our results are in contrast with previous studies showing that triploids tend to be less tolerant to hypoxia than diploids across various fishes (reviewed by Barahona et al., 2026). In particular, Scott et al. (2015) consistently found, across 3 years and 5 rainbow trout strains, that triploid juveniles were less tolerant to hypoxia than diploid counterparts in acute challenge tests with temperature kept at 12°C. These results may be associated to the lower total blood hemoglobin concentrations generally observed in triploids compared to diploid salmonids (Benfey, 1999; Sadler et al., 2000). As reviewed by Benfey (1999), the numbers of red blood cells in triploid fish are fewer than in diploids, but these cells are about 1.4 times as large as their diploid counterparts, which results in triploids having about one-third fewer cells in blood circulation. However, regardless of the strains, Scott et al. (2015) did not see this effect in adult lake-reared trout. Benfey and Devlin (2018) also did not found any significant effect of ploidy on hypoxia tolerance when diploid and triploid rainbow trout were hold for 1 h to temperatures above their thermal optima. We have no clear explanation for our divergent result showing a better hypoxia tolerance of triploid rainbow trout juveniles. We hypothesize that some favourable maternal inbreeding effects in accordance with a 2:1 maternal:paternal gene dosage may be part of the biological explanation as we observed the double amount of phenotypic variance of hypoxia tolerance explained by maternal effects in diploids compared to triploids.

### 4.3. Effect of ploidy on phenotypic and genetic variances, and heritability of traits

In the current study, phenotypic variances of diploid traits were significantly higher than those of triploid performances (with the only exception of FL). This was an unexpected result as higher variances have been generally reported for triploid salmonids (Blanc et al., 2001; Johnson et al., 2007). We hypothesize that the increased phenotypic variance in triploids may partly be explained by developmental instability and chromosome mosaicism due to imperfect physical treatments during the induction process in early studies (Delaval et al., 2024; Glover et al., 2020). However, an alternative explanation has been frequently suggested linked to regulation in gene expressions in response to triploidisation may occur through gene dosage effect or dosage compensation. Dosage effects occur when gene expression correlates with the number of copies of the genes, while dosage compensation occurs when genes are expressed at diploid levels in triploid genomes. An increase in gene dosage is expected to cause increases in the observed phenotypic variance due to elevated additive genetic variance if there is a linear relationship between the level of gene transcription and its effect on phenotype. As triploids often exhibited significantly higher between-family phenotypic variance than diploids for growth traits, it has been concluded that an additive positive dosage effect of gene expressions for growth-related traits is likely in Chinook salmon (Johnson et al., 2007; Shrimpton et al., 2012) as well as in Atlantic salmon full-sib families (Friars et al., 2001; Harvey et al., 2017). However, within-family variance between triploid and diploid full sibs did not always differ significantly between ploidy types (Johnson et al., 2007). Dosage effects, if present, should have a greater impact on the between-family variance component, because the additional set of maternal chromosomes would have a limited effect on within-family variation (as all offspring are more similarly affected), but would exaggerate between maternal family mean trait differences. In general, estimates of maternal variance were larger for triploids than for the diploids (Blanc et al., 2001; Shrimpton et al., 2012), although it has not always been observed (Johnson et al., 2007). However, it should be noticed that when fish are reared in separate family tanks for several months, maternal and environmental effects are confounded. In addition, dominance and/or epistasis effects, if they exist, may inflate the estimates of variances as they could not be disentangled in the studied designs. Unlike diploids, dominance effects may be transmitted across generations in autopolyploids, due to the phenomenon of double reduction. As stated by Hamilton and Kerr (2018), the presence of dominance and epistatic effects would result in overestimation of additive genetic effects and bias in breeding values computed from the application of the **A**-inverse rules.

Our results, based on a mixed-families rearing design, showed that using an incorrect pedigree-based model for triploid expressions lead to large overestimation of genetic variances and heritability values for triploid expressions of production traits, highlighting the importance of accounting for the correct ploidy levels of offspring and parental contributions in the genetic evaluation model. With a proper account of these contributions, pedigree-based and genome-based models yielded similar results regarding heritability estimates and phenotypic variances for all production traits. In addition, we found quite similar heritability estimates for hypoxia tolerance in diploids and triploids under the genome-based model that we considered as the most appropriate model to disentangle additive genetic and maternal effects. The heritability estimates of triploid expressions were also quite close to the estimates for the corresponding diploid expressions for all production traits, at the only exception of SubcutF. for which a lesser heritability was estimated for triploids. This latter result may relate to the fact that body fat is determined during the sexual maturation that does not occur in triploid individuals.

#### Correlation between diploid and triploid traits and expected selection responses on triploid performances

Traits of primary economic interest for fish breeding programs are growth, carcass yield, fillet yield and fat content in muscle (Blay et al., 2021; Haffray et al., 2013) followed nowadays by genetic resistance to pathogens (Houston et al., 2020). Using pedigree information and diploid performances, two breeding programs reported annual genetic gain in the range of 0.5–1% for these three traits as predicted by microwaves or ultrasound technology (Quillet et al., 2005; Vandeputte et al., 2019). The question whether a breeding program may benefit from inclusion of performance data from triploid phenotypes have mainly considered at ‘pan-size’, i.e. about 300 g of weight. Bonnet et al. (1999) observed a significant family × ploidy interaction on body weight, length or condition factor when the fish were about 300g, but not at ‘fillet-size’, about 1 kg. They concluded that selection for improved performance of triploids should be performed at a body weight close to the market size as, while selection based on diploid performance would be effective for ‘fillet-size’, it would be less effective for improvement of ‘pan-size’ triploids. Weber et al. (2014) confirmed these results with low correlations between diploid and triploid body weights when the fish were at or below 300 g and high correlations when the fish were at or above 1 kg.

In the current study, genetic correlations were positive for all traits, signifying that selection for any trait measured on diploids would induce a response in the same direction on the corresponding triploid sibs. These results confirmed previous ones (Leeds and Weber, 2019). In particular, fat-related traits exhibited genetic correlations over 0.95 between diploids and triploids, meaning that very similar genetic responses would be observed in diploid and triploid sibs. For some traits, a slightly higher correlation was estimated between diploid and triploid performances under the correct pedigree-based model and, even more under the genomic model. Although the correlation for all traits were positive and high, some were however significantly different from 1, in particular for body weight and length. Hence, as suggested by Leeds and Weber (2019), it could be better to apply a selection test on triploid traits in order to increase the performance of triploids.

Therefore, we derived the gain or loss in accuracy of selection using triploid collaterals phenotyping rather than diploid phenotyping to improve performances at about 1.5 to 2 kg for slaughtered triploid fish. To summarize, selection response on triploid traits was strongly increased by using a triploid instead of a diploid collateral test when the genetic correlation between diploid and triploid performances were lower than 0.70 such as for BW, moderately increased (∼10%) for correlations about 0.90 (as e.g. for hypoxia tolerance) and the gain was almost null for genetic correlations over 0.95 such as for HGCW, the proxy of fillet yield. Identifying the very primary traits of interest is therefore crucial to determine whether diploid or triploid collateral test should be used in the breeding program

#### Alternative strategies to produce triploids

While the most common strategy for producing triploids in salmonids is applying a pressure treatment to the egg just after fertilization to produce induced-triploids (3n_P), an alternative strategy for producing 3N fish is to mate a tetraploid (4n) parent to a diploid (2n) parent (Chourrout et al., 1986). The resulting 3n progeny are designated intercross-triploids (3n_C). Producing triploids by interploidy alleviate inbreeding problems observed in 3n_P (Hansen et al., 2025) as the tetraploids can be crossbred to increase their heterozygosity. Commercial production of intercross-triploid rainbow trout has not occurred due to technical difficulties associated with the tetraploid induction procedure, and also with fertility-related issues of tetraploid-derived gametes with some males having a too large spermatozoa head size to enable a good fertilization rate (Chourrout et al., 1986; Myers and Hershberger, 1991; Weber et al., 2014). An alternative approach suggested by Hansen et al. (2025) would be to establish a triploid founder broodstock through one generation of gynogenesis. The gynogenesis will eliminate the lethal alleles, and by selecting the best performing individuals from the best performing families, one will also partially eliminate recessive deleterious alleles, increasing the fitness of the resulting triploid population (Hansen et al., 2025).

Nevertheless, the continuous improvement of technical procedures supports that tetraploid production may be feasible. Chourrout et al. (1986) and Weber et al. (2014) showed that the 3n_C grew as well or better than 3n_C and 2n rainbow trout. The higher correlations of body weight between 2n and 3n_C families than between 2n and 3n_P families, particularly at body weights above 1kg, supports that selection based on the 2n phenotype would be considerably more efficient for larger 3NC compared with 3NP fish. In addition, the inferior growth of 3n_P compared with 3n_C fish suggests some negative impacts of the triploid-induction process on performance. Further work should assess the relevance of the previous conclusions that 3n_C fish may have production advantages over 3n_P and 2n fish.

In terms of running pedigree or genome-based BLUP evaluation with the relevant mixture of ploidies (2n, 3n and 4n), the approaches followed in the current study will be successful as the rules proposed by Hamilton and Kerr (2018) would hold to derive the **A**-inverse matrix by changing the value of parameter λ, as well as by Bilton et al. (2024) to derive the **G** matrix considering the right levels of ploidy.

## 5. Conclusion

Our pioneering estimates indicate that heritability values are very similar between diploid and triploid production and quality traits as soon as the ploidy levels of fish is properly accounted for in deriving the pedigree or genome-based relationship matrix used in the genetic evaluation. When maternal effects have to be accounted for such as for early weight or hypoxia tolerance, it is recommended to use a genome-based model to disentangle those effects from the genetic additive values of the fish. Doing so, similar estimates of heritability of hypoxia tolerance in diploids and triploids were observed. Our estimates of genetic correlation between diploid and triploid expressions of the same traits suggest that genetic or genomic evaluation on triploid collaterals may be valuable to increase the accuracy of selection of diploid breeders for growth traits. However, a very limited higher improvement can be expected on fillet yield or hypoxia tolerance. Given the strong correlations estimated between diploid and triploid performances for fillet yield as well as for fat-related traits, selection based on diploid information is very efficient to improve slaughter and quality traits in triploid fish. In consequence, the practical limits of modifying the breeding scheme design well as the economic evaluation of the implementation of triploid performance testing should be assessed by breeding companies across traits and farmed species of interest in aquaculture.

## Declaration of generative AI in the writing process

During the preparation of this work, the authors used google translate and chat GPT to improve the clarity and language of the manuscript. After using these tools, the authors reviewed and edited the content as required and take full responsibility for the content of the publication.

## CRediT authorship contribution statement

Julien Roche: Writing – original draft, Investigation, Formal analysis, Visualization.

Lou Buisson: Investigation, Formal analysis, Visualization.

Pierre Patrice and Yoannah François: Data collection and curation.

Alexandre Desgranges: Resources

Jonathan D’Ambrosio: Project administration, Conceptualization, Funding acquisition.

Pierrick Haffray: Conceptualization, Funding acquisition.

Florence Phocas: Writing – original draft, Supervision, Conceptualization, Funding acquisition.

All co-authors contributed to Writing – review and editing.

## Supplementary data

**S**upplementary data to this article can be found online at XXX

## Appendix A. Lambda estimation

The recombination rate was estimated to be 0.48 in average for the 21,047 markers positioned on the Arlee genome assembly (GenBank assembly accession GCA_013265735.3). The recombination rate evolved along the chromosomes as shown in Supplementary figures 1 and 2; with low recombination rates where centromeres are located and higher rates when moving away from the centromere. The acrocentric chromosomes did not display the “U shape” observed for metacentric ones because their centromeres were really close to telomeres.

**Supplementary figure 1:**
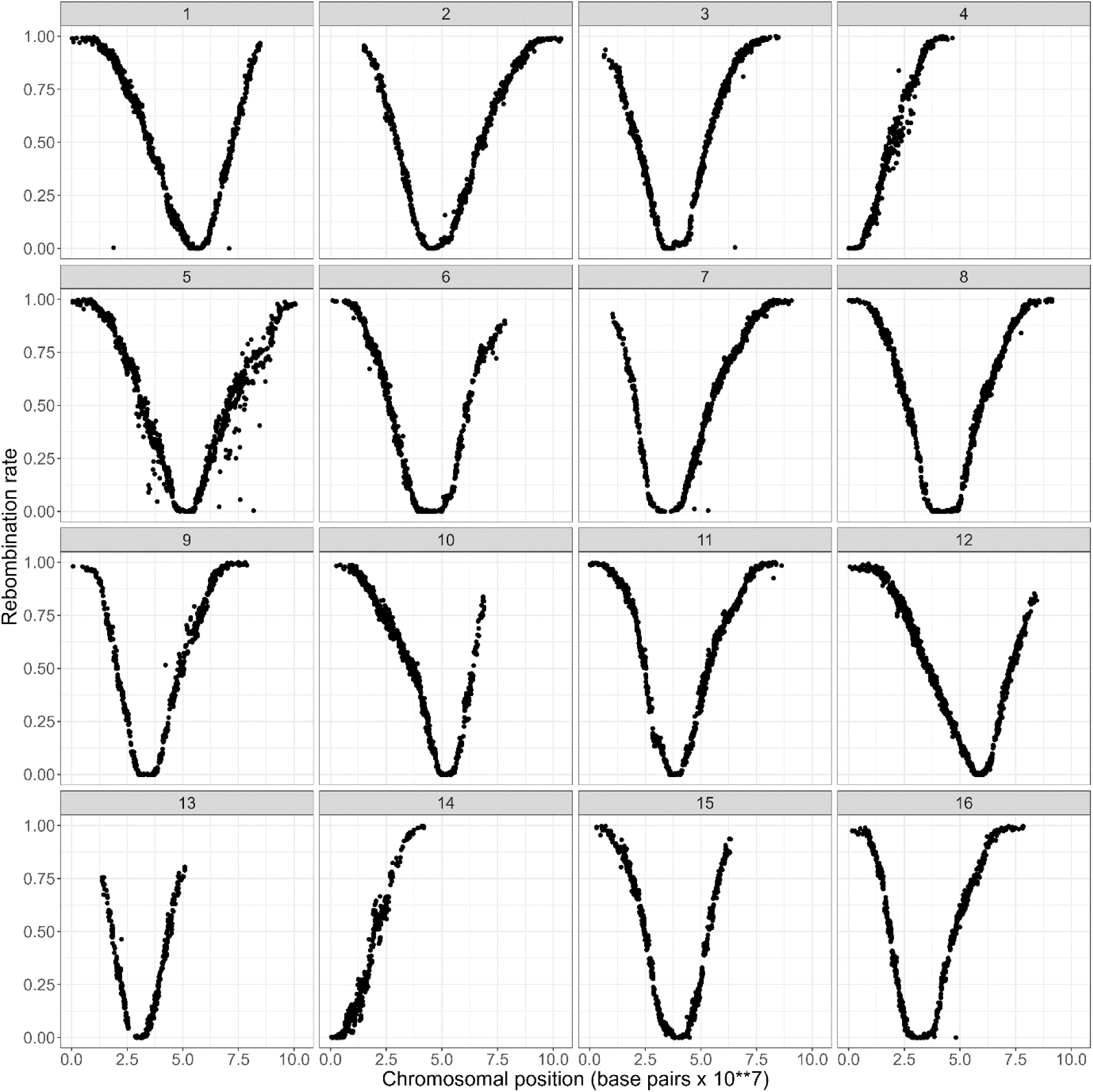
Recombination rate as a function of chromosomal position (base pair) displaying the first 16 chromosomes out of 32 for a total of 12638 markers

**Supplementary figure 2:**
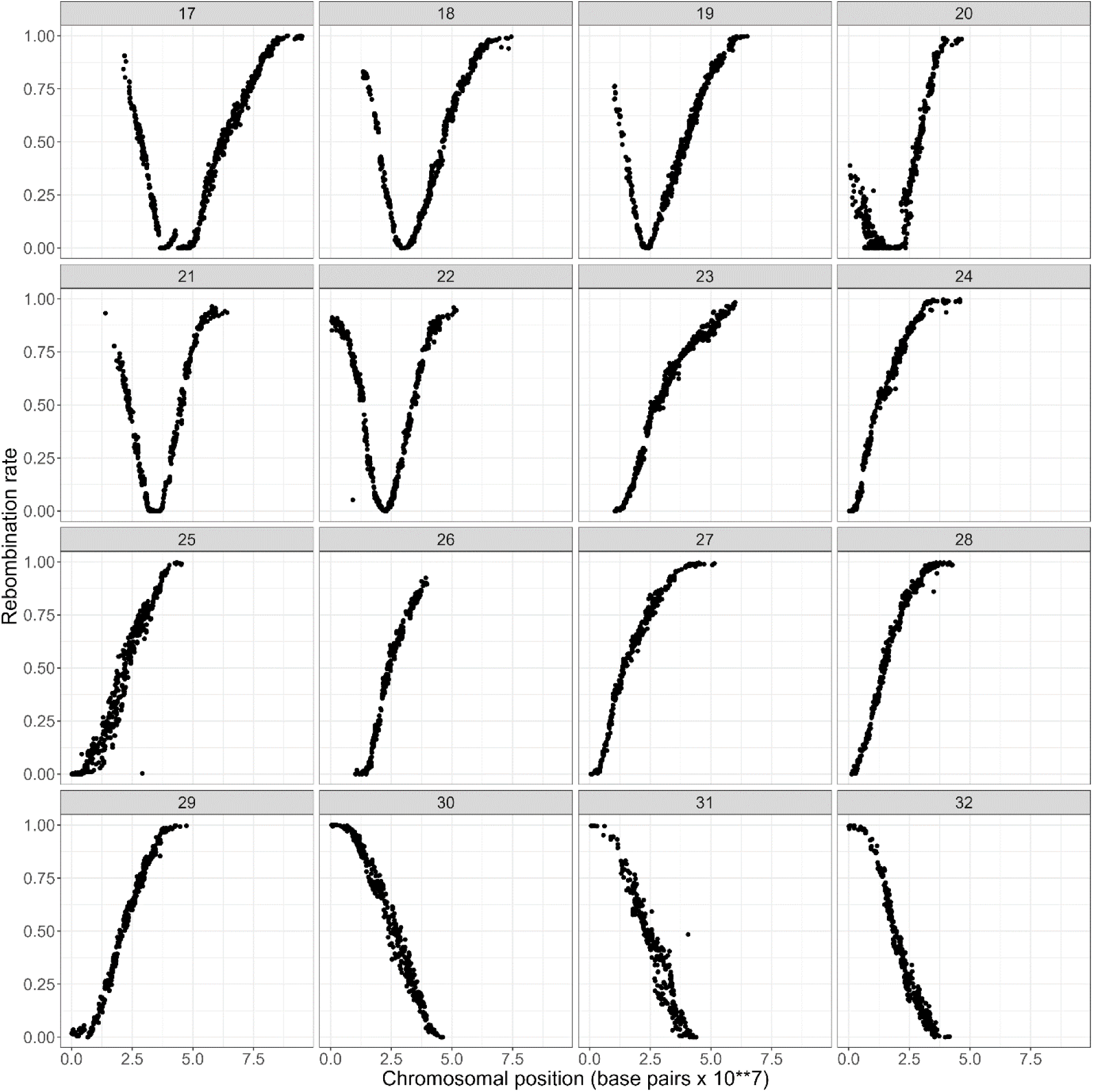
Recombination rate as a function of chromosomal position (base pair) displaying the last 16 chromosomes out of 32 for a total of 7759 markers

