## Supplemental Tables 1 to 4 for "The usefulness of triploid sib records for selection of diploid candidates in rainbow trout breeding programs for growth, processing traits and hypoxia tolerance"

### Supplementary materials

**Supplementary Table 1.** Coefficients (Mean, SD, minimum and maximum values) of the pedigree (A) and genomic (G) relationship matrices used in the genetic evaluation models of production traits, according to the relationship degree and the ploidy level of phenotyped fish.

| **Relationship** | **Ploidy** | **A coefficients** | | | | | **G coefficients** | | | |
| --- | --- | --- | --- | --- | --- | --- | --- | --- | --- | --- |
|  | level | Mean | SD | min | max | Mean | | SD | min | max |
| **Maternal half-sibs** | 2n | 0.277 | 0.032 | 0.250 | 0.383 | 0.240 | | 0.063 | 0.026 | 0.528 |
|  | 3n | 0.683 | 0.020 | 0.667 | 0.771 | 0.636 | | 0.086 | 0.320 | 1.067 |
| **Paternal half-sibs** | 2n | 0.274 | 0.032 | 0.250 | 0.383 | 0.238 | | 0.072 | -0.013 | 0.565 |
|  | 3n | 0.229 | 0.087 | 0.167 | 0.521 | 0.175 | | 0.115 | -0.128 | 0.733 |
| **Full sibs** | 2n | 0.500 | 0.002 | 0.500 | 0.516 | 0.463 | | 0.070 | 0.262 | 0.715 |
|  | 3n | 0.834 | 0.003 | 0.833 | 0.885 | 0.782 | | 0.088 | 0.503 | 1.134 |
| **Dam - offspring** | 2n | 0.501 | 0.003 | 0.500 | 0.516 | 0.464 | | 0.041 | 0.315 | 0.587 |
|  | 3n | 0.817 | 0.004 | 0.816 | 0.848 | 0.778 | | 0.049 | 0.621 | 0.959 |
| **Sire - offspring** | 2n | 0.501 | 0.002 | 0.500 | 0.516 | 0.463 | | 0.041 | 0.259 | 0.600 |
|  | 3n | 0.409 | 0.005 | 0.408 | 0.472 | 0.362 | | 0.050 | 0.194 | 0.542 |
| **Diagonal terms** (1+inbreeding coef.) | 2n | 1.001 | 0.002 | 1.000 | 1.016 | 0.974 | | 0.049 | 0.674 | 1.171 |
|  | 3n | 1.348 | 0.004 | 1.347 | 1.399 | 1.310 | | 0.074 | 1.103 | 1.628 |

**Supplementary Table 2.** Genetic parameters of the production traits estimated by the incorrect pedigree-based model ignoring Hamilton and Kerr’ rules.

| **Traits** | **2n-3n** | **Incorrect diploid parameters** | | | **Incorrect triploid parameters** | | |
| --- | --- | --- | --- | --- | --- | --- | --- |
|  | **r_g_** | **σ²_a2_** | **σ²_P2_** | **h^2^** | **σ²_a3_** | **σ²_P3_** | **h^2^** |
| *Body Weight (BW, g)* | 0.69±0.124 | 33276±7362 | 132799 | 0.25±0.05 | 33519±6054 | 79795 | 0.42±0.06 |
| *Fork Length (FL, mm)* | 0.77±0.151 | 124.2±33.01 | 662.7 | 0.19±0.05 | 213.5±44.94 | 675.8 | 0.32±0.06 |
| *Head Weight (HeadW, g)* | 0.70±0.126 | 336.0±78.28 | 1528 | 0.22±0.05 | 389.6±70.62 | 953.6 | 0.41±0.06 |
| *Viscera Weight (ViscW, g)* | 0.80±0.070 | 706.7±119.0 | 1549 | 0.46±0.06 | 642.7±91.98 | 920.5 | 0.70±0.07 |
| *Gutted carcass weight (GCW*, g)* | 0.87±0.049 | 426.1±64.74 | 735.4 | 0.58±0.07 | 328.6±42.04 | 373.7 | 0.88±0.07 |
| *Headless GCW (HGCW*, g)* | 0.96±0.048 | 450.9±74.14 | 1031 | 0.44±0.06 | 372.2±51.53 | 527.0 | 0.71±0.07 |
| *Muscle Fat (MuscleF, %)* | 0.98±0.029 | 5.817±0.8005 | 8.836 | 0.66±0.07 | 6.349±0.8083 | 7.341 | 0.86±0.07 |
| *Flesh fat (FleshF, %)* | 0.97±0.092 | 2.509±0.3588 | 3.788 | 0.66±0.07 | 2.740±0.3486 | 3.133 | 0.87±0.07 |
| *Subcutaneous fat (SubcutF, %)* | 0.93±0.050 | 5.251±0.8037 | 9.122 | 0.58±0.07 | 5.142±0.7780 | 8.540 | 0.60±0.07 |

* GCW and HGCW are corrected for BW

**Supplementary** **Table 3.** Genetic parameters of the production traits estimated by genomic uni-trait model with only diploid or triploid performances

|  | **Diploid parameters only** | | | **Triploid parameters only** | | |
| --- | --- | --- | --- | --- | --- | --- |
| **Traits** | **σ²_a2_** | **σ²_P2_** | **h^2^** | **σ²_a3_** | **σ²_P3_** | **h^2^** |
| *Body Weight (BW, g)* | 27701±5574 | 132340 | 0.21±0.04 | 18158±3041 | 73882 | 0.25±0.04 |
| *Fork Length (FL, mm)* | 117.4±27.79 | 659.5 | 0.18±0.04 | 123.5±23.58 | 635.5 | 0.19±0.03 |
| *Head Weight (HeadW, g)* | 375.9±70.10 | 1543 | 0.24±0.04 | 247.2±38.64 | 894.9 | 0.28±0.04 |
| *Viscera Weight (ViscW, g)* | 648.5±86.62 | 1542 | 0.42±0.04 | 356.5±44.18 | 808.6 | 0.44±0.04 |
| *Head Weight* (HeadW*, g)* | 95.03±16.48 | 346.8 | 0.27±0.04 | 53.08±8.14 | 188.6 | 0.28±0.04 |
| *Viscera Weight* (ViscW*, g)* | 479.8±48.26 | 730.2 | 0.66±0.04 | 187.7±18.20 | 15.49 | 0.65±0.03 |
| *Gutted carcass weight (GCW*, g)* | 474.8±50.22 | 771.4 | 0.61±0.04 | 181.7±18.44 | 301.9 | 0.60±0.04 |
| *Headless GCW (HGCW*, g)* | 535.1±65.10 | 1065 | 0.50±0.04 | 198.3±23.82 | 447.2 | 0.44±0.04 |
| *Muscle Fat (MuscleF, %)* | 4.651±0.5222 | 8.495 | 0.55±0.04 | 3.176±0.3462 | 6.016 | 0.52±0.04 |
| *Flesh fat (FleshF, %)* | 2.273±0.2373 | 3.681 | 0.62±0.04 | 1.366±0.1488 | 2.561 | 0.53±0.04 |
| *Subcutaneous fat (SubcutF, %)* | 4.253 ±0.5081 | 8.771 | 0.48±0.04 | 3.286 ±0.4191 | 7.699 | 0.43±0.04 |

* Corrected for BW

**Supplementary Table 4.** Genetic parameters of the hypoxia challenge traits estimated by univariate pedigree and genome-based models with only diploid or triploid performances

|  | **Pedigree-based estimates** | | | | | **Genome-based estimates** | | | | |
| --- | --- | --- | --- | --- | --- | --- | --- | --- | --- | --- |
| **Diploid Only** | **σ²_a2_** | **σ²_m2_** | **σ²_P2_** | **h^2^** | **c²** | **σ²_a2_** | **σ²_m2_** | **σ²_P2_** | **h^2^** | **c²** |
| *Weight* | 37.40±10.80 | 7.462±3.903 | 129.6 | 0.29±0.07 | 0.058±0.03 | 28.72±5.865 | 8.770±3.231 | 127.1 | 0.23±0.04 | 0.069±0.02 |
| *TLE* | 1324±403.5 | 405.1±173.6 | 4773 | 0.28±0.08 | 0.084±0.03 | 1221±243.8 | 450.2±145.4 | 4787 | 0.26±0.05 | 0.094±0.03 |
| **Triploid Only** | **σ²_a3_** | **σ²_m3_** | **σ²_P3_** | **h^2^** | **c²** | **σ²_a3_** | **σ²_m3_** | **σ²_P3_** | **h^2^** | **c²** |
| *Weight* | 30.39±9.284 | 5.371±5.405 | 96.84 | 0.31±0.10 | 0.055±0.06 | 16.72±4.189 | 12.68±3.792 | 99.75 | 0.17±0.04 | 0.12±0.03 |
| *TLE* | 402.1±196.3 | 235.6±137.4 | 2677 | 0.15±0.07 | 0.088±0.05 | 489.6±124.1 | 173.0±91.31 | 2672 | 0.18±0.04 | 0.065±0.03 |
